# No tiger in *plastic tiger:* Local linguistic context alters the neural representations of individual words

**DOI:** 10.64898/2026.09.23.753808

**Authors:** Simone M. Krogh, Liina Pylkkänen

## Abstract

Language is vastly ambiguous, yet in context our brains resolve ambiguity effortlessly. This is particularly evident at the lexical level: Although most words can convey a wide range of related meanings, humans rapidly and automatically select the one appropriate for the context. We investigated the neural mechanisms underlying this process by focusing on a case where, intuitively, one meaning appears more basic than another: nouns describing animate entities, such as *tiger*, which can also refer to representations of those entities when combined with a material modifier, as in *plastic tiger*. Using magnetoencephalography, we decoded animacy from neural signals and found that even in such cases, our approach revealed no evidence that the tiger in *plastic tiger* is represented as a sentient animal in any stage of processing. Instead, phrase-level animacy—*plastic tiger* as inanimate—was decodable as early as 210 ms post-stimulus onset and persisted for several hundred milliseconds. These findings challenge theories assuming a shift from a basic sense of a word to a more derived one, highlighting instead the role of compositional context in shaping word meanings.

**Highlights:**

- Phrases like *plastic tiger* have conflicting word- and phrase-level animacy features.
- Using MEG, we found clear evidence for phrase-level animacy encoding at 210 ms.
- In contrast, word-level animacy was never detectable for animacy-mismatch phrases.
- Word meanings are thus not stable entities but shaped by compositional context.

## 1 INTRODUCTION

Patience and Fortitude have guarded the entrance of the New York Public Library for more than a century. Despite their rock-solid appearance, the marble lions are unmistakably that—lions. Such an interpretation of *marble lion* as a representation of an animal, rather than a living creature, arises from the material adjective-noun pairing, with the adjective “coercing” a shift in the noun’s basic meaning (Partee, 2010). This phenomenon is incredibly productive (Wisniewski, 1996), ranging from highly conventionalized (*rubber duck, chocolate bunny*) to more unusual combinations (*plastic tiger*, *porcelain lobster*). Representations of animals, henceforth animacy-mismatch phrases, present an intriguing puzzle at the intersection of linguistic composition and conceptual knowledge: How do our brains interpret *striped/wild/Arabian tiger* as a living, breathing animal but *plastic tiger* as an inanimate representation of that same animal? Specifically, how do we reconcile conflicting semantic features associated with individual words (*tiger* as animate, *plastic* as inanimate) when combining them? Here, we traced the neural evolution of animacy in animacy-mismatch phrases, allowing us to investigate how compositional context modulates individual word meanings.

What, then, constitutes a word’s meaning? While the vast range of potential word uses makes the memorization of every discrete sense implausible (Pustejovsky, 1995), proposals from formal semantics vary in the assumed richness of lexical representations. At one end of the scale, lexical representations are construed as underspecified, consisting of abstract semantic templates that are only populated with appropriate features as a result of the specific context in which a word appears (e.g., Bierwisch & Schreuder, 1992; Reyle, 1993; Blutner, 1998, 2004; Carston, 2012; see also Frisson (2009) for a review of psycholinguistic evidence supporting lexical representations as underspecified). Under such an account, *tiger* in *plastic tiger* is never interpreted as animate since the final composed concept is fundamentally inanimate.

Yet, this view is challenged by the frequently observed divergence in neural activity when constructions involve semantic mismatches such as privative adjectives (*fake gun:* “a gun that is not a gun”; Schumacher et al., 2018; Fritz & Baggio, 2020; Law et al., 2026), metonymic relations (*the ham sandwich wants to pay:* “the customer who ordered the ham sandwich wants to pay”; Rapp et al., 2011; Schumacher, 2014; Weiland et al., 2014; Weiland-Breckle & Schumacher, 2017; Piñango et al., 2017; Yurchenko et al., 2020), and complement coercion (*the author began the book:* “the author began writing/reading/editing the book”; Pylkkänen & McElree, 2007; Brennan & Pylkkänen, 2008, 2010; Pylkkänen et al., 2009; Husband et al., 2011; De Almeida et al., 2016; Lai et al., 2017). These results are often interpreted as reflecting repair processes when the composing context forces a shift away from a word’s basic sense. This interpretation aligns well with the psycholinguistic mechanism of suppression (Hogeweg, 2019), wherein contextually irrelevant features of word meanings appear to be initially activated and only subsequently suppressed (e.g., Swinney, 1979; Glucksberg et al., 2001; Rubio Fernandez, 2007). Taken together, these data support accounts assuming richer lexical representations in which features, including those ultimately incompatible with the context in question, are automatically retrieved and only later pruned (e.g., Pustejovsky, 1995; Jackendoff, 1997; Asher, 2011; Del Pinal, 2015; Vicente, 2015; Asher et al., 2017; for a recent review, see Hogeweg & Vicente (2020)). If so, processing a phrase like *plastic tiger* should first activate the animate feature of *tiger* before suppressing it in favor of the inanimacy of the full phrase.

The conflict between word- and phrase-level animacy makes animacy-mismatch phrases an ideal test bed to adjudicate between word meanings as underspecified or rich. Prior investigations using these constructions have not, however, obtained conclusive evidence for either hypothesis. While a Dutch cross-modal sentential priming study investigating animacy-mismatch phrases (albeit mixed with representations of artifacts like *cloth bike* and *felt cigar*) found some evidence for initial activation of word-related semantic features, consistent with the rich lexical representation hypothesis, no suppression was observed at the investigated stimulus onset asynchrony intervals (0 ms and 400 ms; Hogeweg, 2019). Nevertheless, an ERP study focusing exclusively on animacy-mismatch phrases found them to exhibit distinct neural signatures, eliciting an enhanced positivity at 550–750 ms relative to controls (Schumacher, 2013). Crucially, the functional origins of this late positivity are ambiguous: It could reflect suppression of an activated animate feature that is part of a rich lexical representation, or it could reflect late-stage contextual enrichment of an underspecified semantic template.

To discriminate between the two hypotheses, we employed spatiotemporally resolved MEG measurements to directly test the feature constellations of animacy-mismatch phrases (*plastic tiger*) and controls (*Arabian tiger*). In a set of decoding analyses, we assessed temporal and spatial decodability at the phrase-final word *tiger* using classifiers trained on single-word animates and inanimates. By evaluating the classifier’s predictions against both word-level animacy (where *tiger* is animate in both conditions) and phrase-level animacy (where *tiger* is inanimate in *plastic tiger*), we investigated which meaning is prioritized when linguistic composition puts semantic features at odds.

## 2 METHODS

### 2.1 Participants

Thirty-seven native English-speaking adults completed the experimental protocol. All participants were neurologically intact, had normal or corrected-to-normal vision, and were recruited via the university’s SONA platform or word-of-mouth. After excluding eight participants from analysis due to excessive movement, sleepiness, or reporting a panic attack, the final dataset consisted of 29 participants (16 women, 9 men, 4 nonbinary; 20-40 years old, mean age ± SD: 26.2 years ± 5.1 years). The study was approved by the Institutional Review Board (IRB) ethics committee of [redacted] (approval number: redacted]), and participants were compensated for their time.

### 2.2 Experimental design

Our study compared animacy-mismatch phrases with controls. Because the study was nested within a larger project also investigating noun-noun ordering, stimuli followed a [material/nationality] + [location] + [animal] template, resulting in phrases such as *plastic valley tiger* and *Arabian valley tiger* (Figure 1A). Location-animal pairs were held constant across our experimental conditions, thus keeping lexical material maximally identical while triggering a feature mismatch at the head noun of animacy-mismatch phrases due to the initial material modifier. All stimuli were presented using Rapid Serial Visual Presentation in a 300 ms on, 500 ms off-sequence (see the full trial structure in Figure 1B). We used a simple matching task devoid of metalinguistic reflection adapted from previous research (Pegado et al., 2021; Fallon & Pylkkänen, 2024; Flower & Pylkkänen, 2024, 2026a, 2026b; Dunagan et al., 2025; Krogh & Pylkkänen, 2025; Li & Pylkkänen, 2026). While the task could be performed on purely perceptual grounds, it has been shown to reflect various linguistic properties including, importantly, semantic sensitivity (Fallon & Pylkkänen, 2024, Supplementary Materials). The task was straightforward: Following the sequential presentation of the three-word target stimulus, a full three-word task stimulus was flashed for 300 ms, and participants then had to indicate, via button-press, if the target and task stimuli were identical (match trial) or if one word had been replaced (mismatch trial). Replacements were length-matched and came from the same semantic category. Trials were randomly assigned as match or mismatch trials independently of experimental conditions and varied across participants. Phrases were embedded within blocks of similar three-word phrases manipulating noun-noun order and phrase grammaticality; the findings from those experimental manipulations are reported elsewhere.

**Figure 1:**
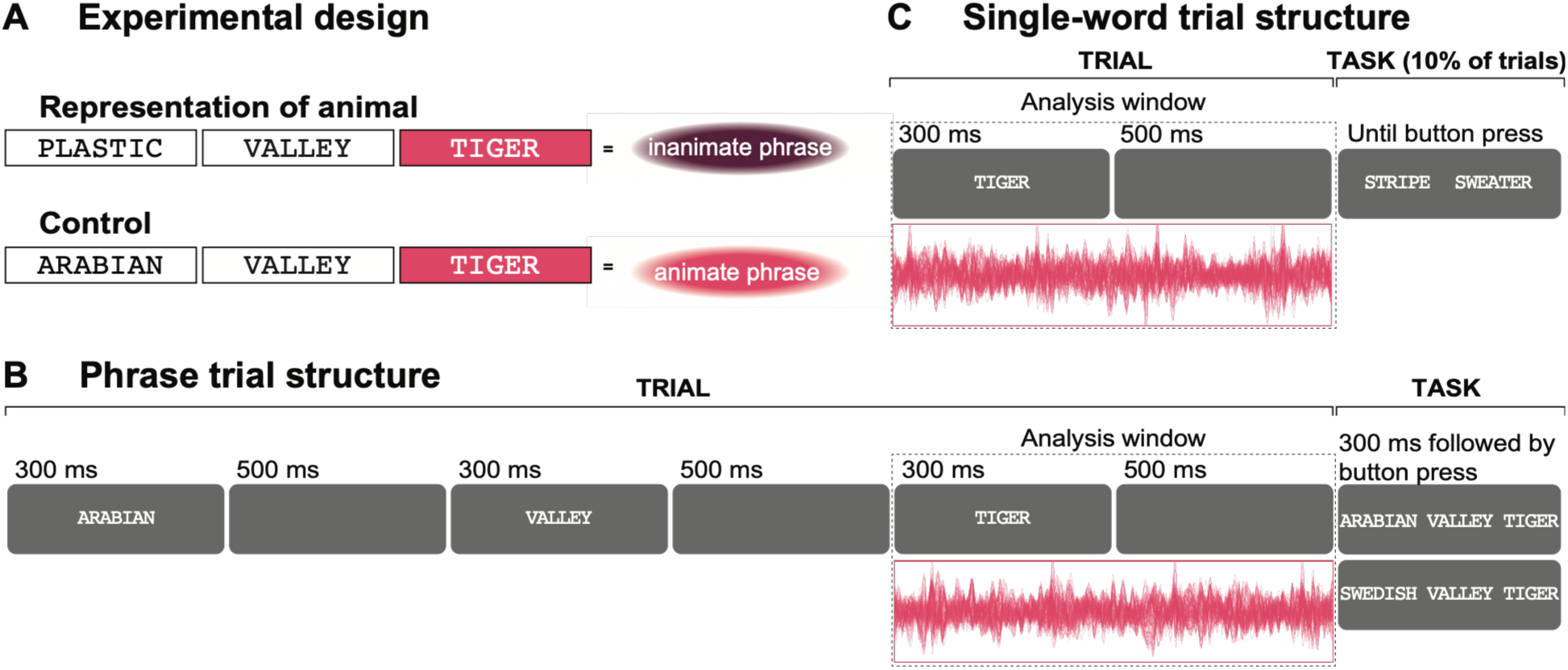
Experimental design and trial structures. (**A**) To examine how the brain prioritizes disjoint semantic features, we contrasted the decodability of word- and phrase-level animacy in animacy-mismatch phrases and controls. Animacy-mismatch phrases denoted inanimate representations of animals, achieved by modifying animate head nouns with a material modifier. Controls were animate at the level of both word and phrase. Location modifiers were included to address a separate question, the results of which are reported elsewhere. (**B**) Phrases were presented word-by-word during continuous MEG recordings and followed by a matching task. The MEG analysis focused exclusively on neural data associated with head nouns (the final word of each phrase). (**C**) Animacy decodability was assessed using single-word animacy as a baseline. Two classifier training sets were constructed, consisting of either the same items used in the phrasal stimuli (ANIMAL/LOCATION) or a mixed group of nouns to test if successful decoding was tied to superordinate semantic categories or extended to animacy as an abstract feature (ANIMATE/INANIMATE). Each word in the classifier training sets was presented individually during continuous MEG recordings, with a semantic association task following 10% of trials.

For our decoding analyses, we trained temporal and spatial classifiers on blocks of single-word trials and assessed their generalizability to head nouns of phrase trials. This cross-condition approach allowed us to determine whether isolated words and words embedded in phrases share stable, discriminable neural patterns associated with animacy. Single-word trials progressed in a similar fashion as phrase trials, with the stimulus being presented for 300 ms and followed by a 500 ms blank screen. To mitigate shallow processing and monitor participant attentiveness, we included a forced-choice semantic association task (e.g., *tiger* → *stripe* | *sweater*) on 10% of trials (Figure 1C). Two types of single-word stimuli were used and presented in separate blocks: the same animals and locations used in the phrases (ANIMAL/LOCATION) and a mixed group of animate and inanimate nouns (ANIMATE/INANIMATE). Due to the considerable length of the full experimental protocol, completion of the ANIMATE/INANIMATE blocks was optional (done by 21 of 29 participants).

### 2.3 Stimuli

#### 2.3.1 Phrases

25 animals and 25 locations matched for lexical characteristics (e.g., length, frequency, semantic, orthographic, and phonological neighbors; Balota et al., 2007; Brysbaert & New, 2009) formed the basis of the phrasal stimuli. First, each animal was modified by two plausible locations (e.g., *tiger* by *valley* and *desert*). These location-animal pairs were then further modified by one of ten constitutive materials (e.g., *plastic*) and ten length-matched geographical place names (e.g., *Arabian*), ensuring that all pairings remained ecologically plausible (i.e., excluding improbable combinations like *Swedish valley tiger*). This yielded 50 animacy-mismatch phrases such as *plastic valley tiger* and 50 matched controls like *Arabian valley tiger*. An additional 100 trials were created in which the location-animal order was swapped (e.g., *Arabian tiger valley*). Regardless of noun-noun order, all location-animal and animal-location combinations were novel (transition probability: 0% in COCA (Davies, 2008); < 0.07 in iWeb (Davies, 2018)). Finally, 200 ungrammatical phrases were created by swapping the first two words of a grammatical phrase (grammatical: *Arabian valley tiger*; ungrammatical: *valley Arabian tiger*). All stimuli were presented in upper-case letters to avoid bias from conventional capitalization of geographical place names.

#### 2.3.2 Single words: Animals and locations

The 25 animals and 25 locations used in the phrases were presented in isolation to serve as a training set for decoding animacy at the level of superordinate semantic categories (ANIMAL/LOCATION). Each exemplar was repeated ten times across ten blocks, resulting in a total of 500 trials (250 trials per condition).

#### 2.3.3 Single words: Animates and inanimates

To test if results obtained using the ANIMAL/LOCATION training set reflect this specific category contrast or animacy encoding more generally, we created an additional classifier training set (ANIMATE/INANIMATE). This set comprised 25 animate and 25 inanimate nouns that spanned different superordinate categories and were matched for the same lexical characteristics as the ANIMAL/LOCATION training set. Inanimate stimuli included tools, vehicles, buildings, musical instruments, and pieces of clothing, whereas animate stimuli all referred to human or human-like individuals such as *tourist, emperor, dentist,* or *vampire.* Note that animal and location exemplars were not included in this ANIMATE/INANIMATE contrast, which unfortunately limited the possible diversity of superordinate categories in the ANIMATE set. Again, each exemplar was repeated ten times across ten blocks, resulting in a total of 500 trials (250 trials per condition).

### 2.4 Procedure

Upon arrival to the [redacted] lab, participants provided informed consent and completed a brief demographic questionnaire. Next, digitizations of participants’ head shapes alongside five future marker coil placements and three anatomical landmarks (nasion as well as left and right preauricular points) were acquired with a Polhemus FastSCAN system. Participants were then led through a practice session spanning all experimental components and subsequently guided into the magnetically shielded room hosting the MEG machine. Participants completed the experimental protocol during continuous MEG recordings while lying down. The positions of marker coils were recorded at the beginning and end of the recording session.

The experimental protocol had three obligatory experimental components: single-word ANIMAL/LOCATION trials (10 blocks of 50 trials), RSVP phrase trials (4 blocks of 100 trials), and a third component flashing the same phrases all-at-once rather than word-by-word (4 blocks of 100 trials). This third component was included to assess whether basic combinatory processing generalizes across presentation modes, and results are reported elsewhere. Blocks from the three obligatory experimental components were interleaved in a semi-structured, pseudorandomized manner with appropriate counterbalancing. Interleaving blocks ensured that data from the three obligatory components were sampled early and late in the recording session to account for potential participant fatigue. On average, it took participants around 70 minutes to complete all 18 blocks. If time allowed it, participants were then given the option to complete the optional single-word ANIMATE/INANIMATE component of approximately 20 minutes. This was entirely voluntary given the considerable length of the core protocol, and participants received additional compensation if opting to complete it. Participants were instructed to take breaks as needed throughout the recording session.

### 2.5 MEG data collection and preprocessing

MEG data were acquired using a whole-head, 157-channel axial gradiometer system (Kanazawa Institute of Technology, Kanazawa, Japan) at a 1000 Hz sampling rate and a 0.1-200 Hz online bandpass filter. A photodiode measured the latency between triggers and visual stimulus onset. Initial noise reduction was performed in the MEG160 software, using the Continuously Adjusted Least-Squares Method (Adachi et al., 2001) to remove environmental noise via three reference channels. All subsequent preprocessing steps were performed in Python using *MNE-Python* (v.1.10; Gramfort et al., 2014).

The data were first bandpass-filtered between 1 and 40 Hz, with the high-pass filter being necessary to attenuate NYC environmental noise. Two known broken channels, along with recording-specific flatlined or excessively noisy channels (mean: 1.45 additional bad channels), were removed and interpolated based on neighboring channels. The data were then decomposed via Independent Component Analysis (ICA) and cleaned of ocular, cardiac, and well-characterized environmental artifacts. Phrase trials were segmented into three separate 800 ms epochs (300 ms word presentation; 500 ms blank screen) and baseline corrected with the 100 ms preceding the first word. We focused exclusively on the epochs associated with the third word. Single-word trials were segmented into one 800 ms epoch (300 ms word presentation; 500 ms blank screen) with a 100 ms pre-stimulus baseline. Epochs were automatically rejected if peak-to-peak amplitude exceeded 3000ft at any point. Additionally, we excluded trials manually if they were flagged during data acquisition, e.g., due to technical issues or if participants exhibited task irrelevant behavior such as talking to the experimenter. Phrase epochs were further cleaned of trials with incorrect responses or response times more than three SDs from the participant mean. On average, 45 epochs (SD = 5.1) were kept for the two phrase conditions analyzed here. Similarly, single-word epochs were cleaned of trials with incorrect responses and trials immediately following the semantic association task (occurring only on 10% of trials) to avoid potential spillover effects of task-induced surprisal. This yielded an average of 222.6 epochs (SD = 5.9) per condition for the ANIMAL/LOCATION single-word trials, and 223.4 epochs (SD = 4.3) per condition for the ANIMATE/INANIMATE single-word trials.

We estimated source-level activity from single-trial evoked responses via Dynamic Statistical Parameter Maps (dSPM; Dale et al., 2000). First, we scaled and coregistered the FreeSurfer *fsaverage* brain (Fischl, 2012) to each participant’s digitized head shape and fiducial markers. The use of a template brain in lieu of individual MRIs thus calls for appropriate caution when interpreting source localized results. On the scaled *fsaverage* surface, we generated a source-space mesh of 2,562 vertices per hemisphere (ico-4 spacing). Then, forward solutions were computed for each participant using the Boundary Element Model (BEM) method alongside channel noise-covariance matrices using the 100 ms pre-stimulus interval of all experimental trials. Based on these, subject-specific inverse solutions were calculated and applied to the single-trial data using an SNR of 3. This yielded L2 minimum norm source estimates that were noise-normalized to produce the final dSPM values.

### 2.6 Statistical analysis

#### 2.6.1 Behavioral data

Mirroring our MEG preprocessing, the behavioral cleaning pipeline excluded trials flagged during data acquisition as well as any phrase trials with response times > 3 SDs from a participant’s mean. For the single-word trials, we simply report mean accuracies. For the phrase trials, we constructed linear and generalized linear mixed-effects models to examine the influence of initial modifier (*plastic, Arabian*) and response type (match trial, mismatch trial). Prior research has shown match and mismatch trials to incur distinct processing demands (Fallon & Pylkkänen, 2024; Flower & Pylkkänen, 2024, 2026a; Krogh & Pylkkänen, 2025; Li & Pylkkänen, 2026), thus meriting its inclusion as a factor in the design. Using accuracy and log-transformed response times (from correct trials only) as dependent variables, we employed the maximal random-effects structure that reached convergence, with intercepts for participants and items. Statistical significance was determined using likelihood ratio tests; where applicable, pairwise comparisons were corrected using a Tukey adjustment. Although we focused our analysis on the two experimental conditions central to this study, the results remain qualitatively consistent when modeled with the full experimental design. All analyses were conducted using the *lme4* (Bates et al., 2015) and *afex* (Singmann et al., 2023) packages in R (v4.5.2) and RStudio (v2026.01).

#### 2.6.2 Temporal decoding analyses with generalization across time

Our primary goal was to trace the neural encoding of word- and phrase-level animacy, which we did using time-resolved decoding with temporal generalization in sensor space (King & Dehaene, 2014). Training classifiers on spatial neural patterns at one time point and testing across all others allowed us to identify shared representations even if they differed in latencies across training and testing sets.

Classifiers were trained on either single-word ANIMAL/LOCATION or ANIMATE/INANIMATE trials and evaluated on the head noun of phrases (e.g., *tiger;* Figure 2). We restricted decoding to the head noun because successful classification at this point can unambiguously be attributed to either lexical or phrasal animacy. In contrast, animacy encoding of the preceding location modifiers (e.g., *valley*) is inherently ambiguous: Successful classification at this stage could reflect word-level animacy (*valley* as inanimate) or anticipatory encoding of the entire phrase-level representation (*plastic valley tiger* as inanimate). Initial modifiers were excluded from analysis as geographical adjectives are underspecified for word-level animacy, and phrase-level animacy could only be determined upon seeing the second word.

**Figure 2:**
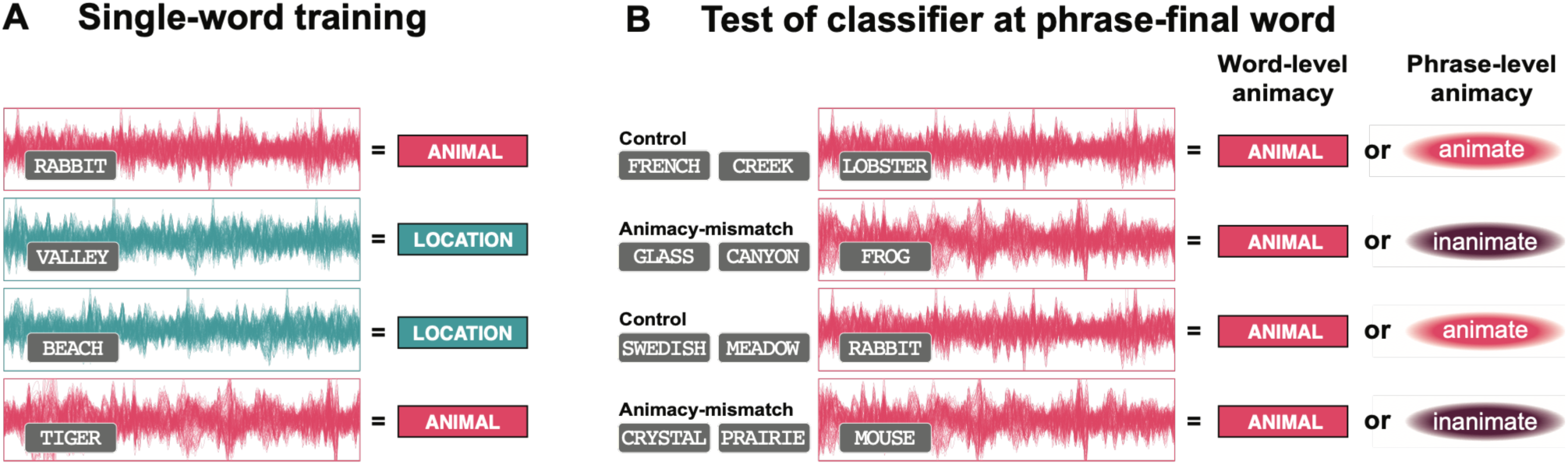
Decoding analysis pipeline. (**A**) We trained classifiers on the neural data of single-word trials alongside appropriate category labels (e.g., ANIMAL vs. LOCATION). (**B**) Each classifier was tested on head nouns in phrases using two labeling schemes, allowing us to disentangle word-level from phrase-level animacy encoding. Under the word-level labeling scheme, all trials were assigned the “animate” label. Under the phrase-level labeling scheme, controls retained the “animate” label while animacy-mismatch phrases were assigned the “inanimate” label. This basic pipeline was used for our temporal and, where applicable, spatial decoding analyses.

To determine if neural data at the head noun contained reflexes of animacy at the word-level, phrase-level, or both, we implemented two distinct labeling schemes. First, we tested our single-word-trained classifiers against head-noun data using a word-level animacy labeling scheme, where all instances of *tiger,* e.g., in *plastic valley tiger* and *Arabian valley tiger,* had the label “animate”. Second, we repeated the process but assigned trial labels using a phrase-level animacy labeling scheme. While *tiger* in *Arabian valley tiger* thus retained its “animate” label, *tiger* in *plastic valley tiger* was relabeled “inanimate”. Crossing our two training sets (ANIMAL/LOCATION, ANIMATE/ INANIMATE) with the two labeling schemes (word-level, phrase-level) yielded four independent decoding analyses.

The details of our temporal decoding pipeline were as follows. For both training and test sets, the MEG data were downsampled to 200 Hz by averaging non-overlapping bins of 5 ms. The data dimensionality of training sets was then reduced from 157 sensors to 70 principal components, accounting for at least 96.6% of the variance. This PCA mapping was subsequently applied to the test data. All data were scaled to unit variance and fed to a logistic regression classifier with l2 regularization. Regularization strength (C) was optimized using a grid search on a logarithmic scale (10^-4^ to 10^4^) with stratified 5-fold cross-validation on the training set. Separate classifiers were trained on pairs of time points in windows of 100 ms (with edge padding) and repeated for every 10 ms step throughout the 0–800 ms epoch. This produced a temporal generalization matrix for each analysis. The entire pipeline was done separately for each participant.

#### 2.6.3 Group-level statistical testing of temporal decoding analyses

Group-level temporal decoding accuracy was evaluated against a chance baseline of 0.5 using non-parametric cluster-based permutation tests (Maris & Oostenveld, 2007). At each train-test time pair, we conducted one-tailed, one-sample *t* tests, aggregating contiguous *t* values (*p* < 0.05) into clusters with a minimum duration of 20 ms. To determine cluster-level significance, we compared observed cluster statistics against a surrogate null distribution generated via 10,000 within-subject label permutations. Observed clusters were assigned corrected *p* values based on their frequency relative to this surrogate distribution. In accordance with Sassenhagen & Draschkow (2019), cluster boundaries are reported as approximations as they originate from uncorrected *t* values.

Following standard practice, we initially performed group-level analyses by averaging participant accuracies across the two experimental conditions. This aggregate approach identified clusters where the neural encoding of *tiger* converged across *plastic valley tiger* and *Arabian valley tiger* under a specific labeling scheme (word-level animacy, phrase-level animacy). Importantly, aggregate clusters do not necessarily imply uniform contributions across experimental conditions. A cluster may be disproportionately driven by a single condition; conversely, robust effects in one condition may be neutralized by poor performance in the other. To address this, we also conducted condition-specific analyses for *plastic valley tiger* and *Arabian valley tiger,* respectively. It should be noted that these condition-specific analyses necessarily had lower trial counts than the aggregate analyses, which likely reduced statistical power.

We report results from three train-test windows: 100–400 ms, 400–700 ms, and 100–700 ms. The latter analysis window represents the maximum viable temporal range within our 0–800 ms epoch, accounting for the 100 ms edge padding imposed by the sliding window architecture. To capture effects that may not survive cluster permutations in this considerably broad train-test window, we included the 100–400 ms train-test window to capture lexical access and the 400–700 ms train-test window to target the late positivity previously associated with animacy-mismatch phrases (Schumacher, 2013). MEG analysis scripts were written in Cursor, an AI-assisted code editor used for limited code writing, editing, and debugging with both automatic model routing (Auto) and manually selected Claude Sonnet and Opus models (4.5 and 4.6). All aspects of the decoding analyses were performed in Python using *MNE-Python* (v1.0.3) and *scikit-learn* v1.6.1 (Pedregosa et al., 2011).

#### 2.6.4 Spatial decoding analyses

Despite the multivariate nature of temporal decoding, some brain regions may contribute more to differences in spatial neural patterns than others. We therefore followed our temporal decoding analyses with targeted spatial decoding analyses in significant temporal clusters. This involved fitting unique classifiers to temporal neural patterns using a 10mm-radius searchlight centered at each individual neural source. Implementation details are otherwise as described in Section 2.6.2. To systematically evaluate the full extent of significant clusters, each significant cluster was subdivided into 50 ms × 50 ms train-test tiles for separate spatial decoding analyses. Tile grids were uniformly aligned across temporal clusters of the same analysis type (e.g., condition-specific for animacy-mismatch phrases) arising from the two training sets (ANIMAL/LOCATION, ANIMATE/INANIMATE), positioned such that the window of maximal temporal overlap fell on a tile diagonal. We excluded any tiles where significant temporal cluster points constituted less than 20% of the total tile area. Again, the procedure was done separately for each analysis and each participant.

#### 2.6.5 Group-level statistical testing of spatial decoding analyses

Group-level significance within each temporal tile was assessed using cluster-based permutation tests (see Section 2.6.3) in predefined regions of interests (ROI). Within each temporal tile, *p*-values were corrected for multiple comparisons across ipsilateral ROIs using the false discovery rate (FDR) procedure (Benjamini & Hochberg, 1995) at an adjusted significance threshold of *p* < 0.05. Because these spatial decoding analyses targeted previously identified temporal clusters, they are post-hoc in nature and the corrected *p* values should be viewed as descriptive of spatial localization rather than independent inferential statistics.

We selected four ROIs based on two criteria. First, since animacy-mismatch phrases have not previously been investigated using spatially resolved neuroimaging techniques, we identified regions previously implicated in the processing of semantically complex constructions, such as metonymy and coercion. This yielded three ROIs: ventromedial prefrontal cortex (vmPFC; Pylkkänen & McElree, 2007; Brennan & Pylkkänen, 2008, 2010; Pylkkänen et al., 2009; De Almeida et al., 2016; Piñango et al., 2017), the inferior frontal gyrus (IFG; Husband et al., 2011; Rapp et al., 2011; De Almeida et al., 2016; Lai et al., 2017), and posterior parts of the temporal lobe including the superior temporal gyrus and the superior temporal sulcus (PTL; De Almeida et al., 2016; Rapp et al., 2011). However, because these studies employed difference-rather than similarity-based methodologies like our decoding approach, we also sought to identify regions likely to be sensitive to shared conceptual representations. While the vmPFC is one such candidate, given its hypothesized role in representing the output of combinatory processes (Pylkkänen, 2019), we likewise targeted the anterior temporal lobe (ATL) based on its well-established role as a semantic hub (Lambon Ralph et al., 2006; Ralph et al., 2017) and implication in conceptual composition (Bemis & Pylkkänen, 2011; Westerlund & Py-lkkänen, 2014; Zhang & Pylkkänen, 2015). The vmPFC was defined as the combination of BAs 10/11 and the IFG as BAs 44/45. The ATL and PTL were defined using the *aparc* atlas (Desikan et al., 2006), available with Freesurfer. While our focus was on the left hemisphere ROIs, we included the right hemisphere homologues because some previous studies found bilateral or right-lateral effects in the identified ROIs.

## 3 RESULTS

### 3.1 Behavioral processing costs for conflicting animacy features in mismatch trials

Our linear mixed-effects regression model for response times yielded significant main effects of initial modifier (*p* = 0.011) and response type (*p* < 0.001), along with a robust interaction between the two (*p* < 0.001; Figure 3 and Table 1). Pairwise comparisons clarified that this interaction was driven by a processing cost for animacy-mismatch phrases in mismatch trials (*p* < 0.0001; animacy-mismatch phrases: 865.74 ± 386.7 ms; control: 839.39 ± 383.01 ms). We saw no significant difference in accuracy between experimental conditions, which elicited high accuracies across-the-board (animacy-mismatch phrases: 93.53 ± 6.7%; controls: 94.36 ± 6.11%). As in prior studies (Fallon & Pylkkänen, 2024; Flower & Pylkkänen, 2024, 2026a; Krogh & Pylkkänen, 2025; Li & Pylkkänen, 2026), match trials enjoyed a processing advantage over mismatch trials (*p* < 0.001), but this did not interact with initial modifier type (*p* = 0.769). Mean accuracies for each of the semantic association tasks were high (ANIMAL/LOCATION: 95.01 ± 21.78%; ANIMATE/INANIMATE: 95.33 ± 21.11%).

**Figure 3:**
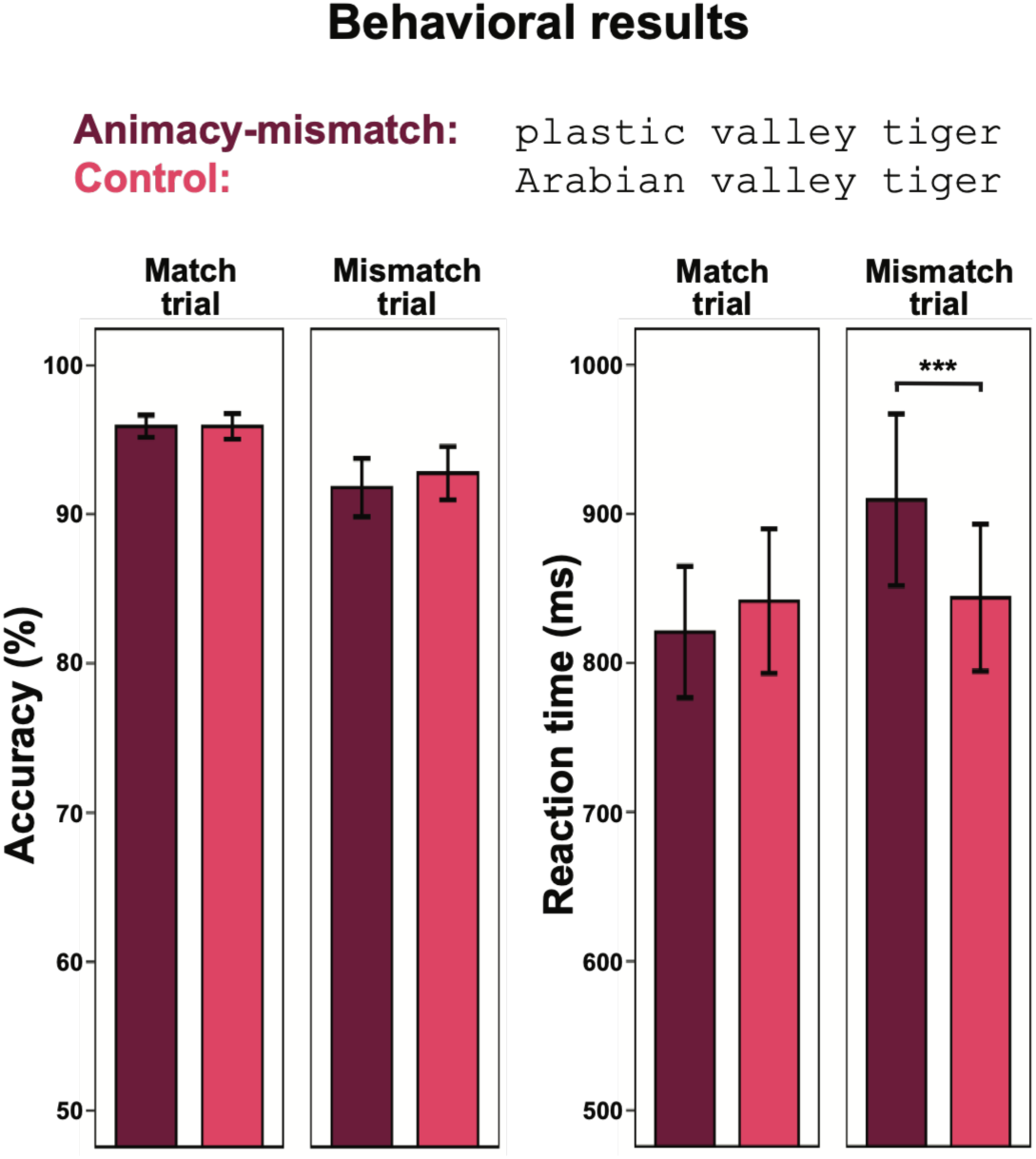
Behavioral results across reaction time and accuracy. Animacy-mismatch phrases and controls were mostly processed identically, although animacy-mismatch phrases in mismatch trials had longer response times relative to controls.

**Table 1:** Likelihood ratio tests of behavioral data. Log-transformed reaction times (incorrect responses excluded) were modeled using linear-mixed effects regression while accuracy was modeled using generalized linear mixed-effects regression.

| **Effect** | **df** | **Reaction time** |  | **Accuracy** |  |
| --- | --- | --- | --- | --- | --- |
|  |  | **$\chi^2$** | ***p* value** | **$\chi^2$** | ***p* value** |
| initial modifier | 1 | 6.43 | 0.011 | 0.46 | 0.497 |
| response type | 1 | 34.87 | < 0.001 | 16.78 | < 0.001 |
| initial modifier × response type | 1 | 12.95 | < 0.001 | 0.09 | 0.769 |

### 3.2 Distributed neural patterns encode phrase-level, not word-level, animacy

Our temporal decoding analyses with generalization over time employed two distinct labelling schemes, allowing us to compare the neural encoding of word- and phrase-level animacy. This labeling switch specifically targeted animacy-mismatch phrases (*plastic valley tiger*), where a conflict exists between the animacy of the head noun and the full phrase. As described in *Group-level statistical testing of temporal decoding analyses,* we evaluated group-level performance using both an aggregate approach (averaging across both experimental conditions) and a condition-specific approach (averaging each experimental condition independently).

Temporal decoding was successful only when classifying words based on phrase-level, and not word-level, animacy. Within the 100–700 ms time window, the classifier trained on single-word ANIMAL/LOCATION trials and tested using the phrase-level animacy labeling scheme revealed two significant off-diagonal clusters (Figure 4A). First, an early aggregate cluster emerged with train times ∼100 ms to ∼310 ms and test times ∼210 ms to ∼400 ms (*p* = 0.0446). Later, a prolonged cluster specific to animacy-mismatch phrases appeared at train times ∼260 ms to ∼380 ms and test times ∼ 420 ms to ∼700 ms (*p* = 0.046). In both cases, test times were delayed relative to train times. In the 100–400 ms window, we additionally observed a marginal diagonal cluster specific to controls with train times ∼200 ms to ∼310 ms and test times ∼190 ms to ∼350 ms (p = 0.0793). The 400–700 ms window did not yield any significant clusters.

**Figure 4:**
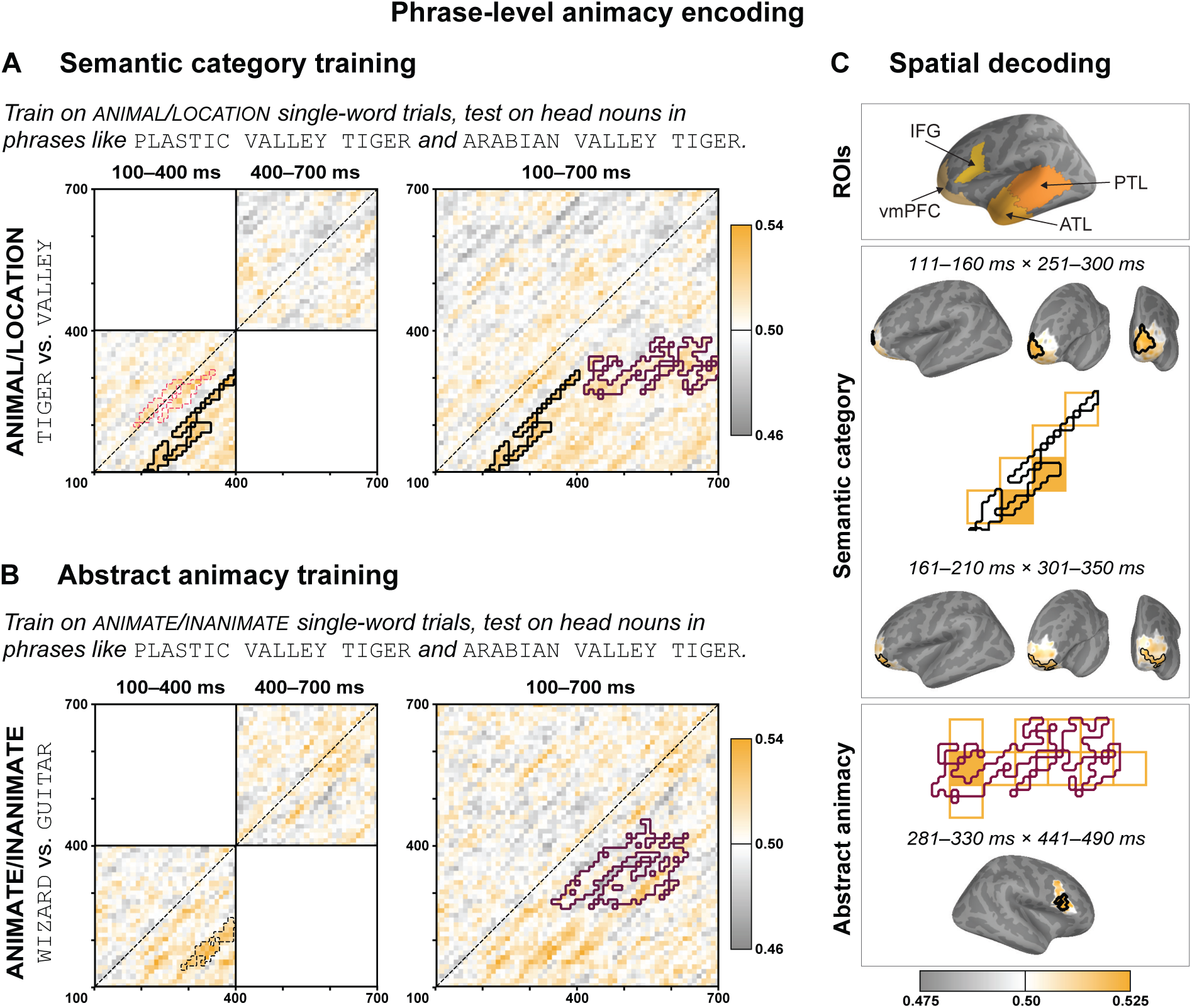
Phrase-level animacy encoding. (**A**) Two significant clusters emerged for classifiers trained on ANIMAL/LOCATION single-word trials and tested on head nouns using *the phrase-level labeling scheme*. An early cluster (train: ∼100–310 ms; test: ∼210–400 ms) arose from the aggregate group-level analysis in the 100–400 ms and 100–700 ms time windows, providing evidence of early representation of compositional output. The second significant cluster arose from the condition-specific analysis for animacy-mismatch in the 100–700 ms time window, characterized by sustained generalization from training to test data (train: ∼260–380 ms; test: ∼420–700 ms). Finally, the condition-specific analysis for controls yielded a marginal cluster on the diagonal in the 100–400 ms window. Note that the different trial counts in the aggregate and condition-specific analyses hinder direct comparisons. (**B**) Classifiers trained on ANIMATE/INANIMATE single-word trials (completed by 21 of 29 participants) produced similar results, with a trending, early aggregate cluster (train: ∼140–240 ms; test: ∼290–400 ms) and a significant, sustained cluster from the condition-specific analysis for animacy-mismatch phrases (train: ∼270– 450 ms; test: ∼350–630 ms). Heatmaps in (**A**) and (**B**) reflect decoding accuracies of the aggregate group-level analyses. (**C**) Spatial decoding localized parts of the effects obtained for classifiers trained on ANIMAL/LOCATION single-word trials, not for those trained on ANIMATE/INANIMATE single-word trials. The aggregate cluster localized to the left ventromedial prefrontal cortex while the animacy-mismatch cluster localized to the right inferior frontal gyrus. vmPFC: ventromedial prefrontal cortex; IFG: inferior frontal gyrus; ATL: anterior tem-poral lobe; PTL: posterior temporal lobe. Solid outline: *p* < 0.05; dashed outline: 0.05 < *p* < 0.1. Outline colors: black = aggregate cluster; burgundy = animacy-mismatch cluster; magenta = control cluster.

Turning next to our ANIMATE/INANIMATE training set, which were completed by a subsample of participants (21 of 29), we saw similar patterns when using the phrase-level animacy labeling scheme (Figure 4B). This included a trending, early off-diagonal aggregate cluster with train times ∼140 ms to ∼240 ms and test times ∼290 ms to ∼400 ms (*p* = 0.0665) in the 100–400 ms window followed by a significant, prolonged off-diagonal cluster specific to animacy-mismatch phrases with train times ∼270 ms to ∼450 ms and test times ∼350 ms to ∼630 ms (*p* = 0.0229) in the 100–700 ms window. Again, the 400– 700 ms window did not yield any significant clusters.

Our word-level animacy labeling scheme reproduced the early trending cluster for our controls but did not produce any new clusters (Figure 5).

**Figure 5:**
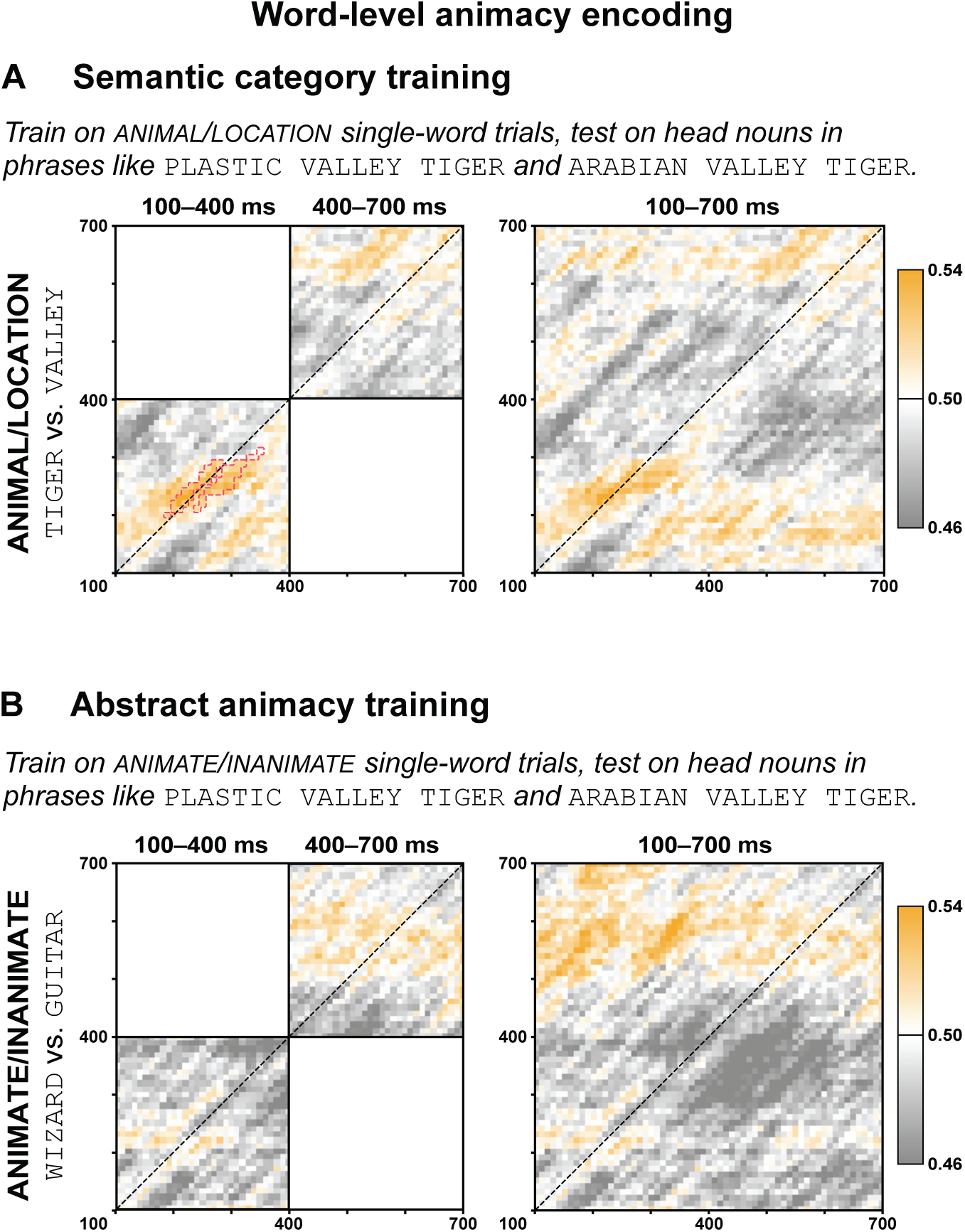
Word-level animacy encoding. (**A**) Classifiers trained on ANIMAL/LOCATION single-word trials and tested on head nouns using the word-level labeling scheme reproduced the marginal cluster for controls but yielded neither aggregate nor animacy-mismatch clusters. (**B**) No clusters arose from training on ANIMATE/INANIMATE single-word trials under the word-level labeling scheme. Solid outline: *p* < 0.05; dashed outline: 0.05 < *p* < 0.1. Outline colors: black = aggregate cluster; burgundy = animacy-mismatch cluster; magenta = control cluster.

### 3.3 Left vmPFC and right IFG involvement in phrase-level animacy encoding

We localized the significant temporal decoding effects using spatial decoding analyses, subdividing temporal clusters into 50 ms × 50 ms train-test tiles. When probing the aggregate cluster obtained from training on single-word ANIMAL/LOCATION trials, we observed a significant cluster in the left vmPFC in the 111–160 ms × 251–300 ms tile (corr. *p* = 0.0332) as well as a trending left vmPFC cluster in the ensuing 161–210 ms × 301–350 ms tile (corr. *p* = 0.0692). No tiles in the aggregate cluster trained on single-word ANIMATE/INANIMATE trials yielded significant clusters. Turning next to the clusters specific to animacy-mismatch phrases trained on single-word ANIMAL/LOCATION trials, a single cluster emerged in the right IFG in the 281–330 ms × 441–490 ms tile (corr. *p* = 0.0372; Figure 2.4C). Again, no clusters emerged when training on single-word ANIMATE/INANIMATE trials.

## 4 DISCUSSION

In this work, we investigated how the brain prioritizes competing semantic features during linguistic composition, offering insights into the architecture of lexical representations. Our experimental conditions juxtaposed animacy-mismatch phrases (*plastic tiger*), where word-level animacy of the head noun dissociates from that of the phrase, with controls (*Arabian tiger*). In a series of decoding analyses, we isolated neural reflexes of word-level animacy from those of the phrase by training on single words differing in animacy (ANIMAL/LOCATION; ANIMATE/INANIMATE) and testing the classifier’s ability to generalize learned representations to the head noun under different labeling schemes. This allowed us to test whether word-level animacy is encoded even when contextually irrelevant, or if phrase-level animacy takes precedence. Under the former hypothesis, lexical representations would be conceptually rich by default and subsequently pared down to fit a given compositional structure. Alternatively, the presence of phrase-level animacy encoding in the absence of word-level effects would point to lexical representations as sparse semantic templates that undergo conceptual specification based on the context in which they appear.

Our results show that the brain prioritizes phrase-level over word-level meanings, thus supporting the hypothesis positing lexical representations to be enriched by context. While we found no evidence of word-level animacy encoding for our animacy-mismatch phrases—the condition allowing us to disentangle lexical and phrasal features—phrase-level animacy was decodable from distributed neural patterns in two stages. In the first stage, neural patterns associated with the processing of animals and locations in isolation (ANIMAL/LOCATION) between ∼100–310 ms mapped to phrase-level animacy representations at ∼210–400 ms. This effect emerged from the combined activity across experimental conditions, with spatial decoding analyses revealing the left vmPFC as one locus of activity. The second stage was specific to animacy-mismatch phrases, characterized by a prolonged mapping of single-word animacy features at ∼260–380 ms onto phrase-level representations at ∼420–700 ms. Spatial decoding localized this sustained effect in part to the right IFG. Across-the-board, similarly timed phrase-level animacy effects emerged when animacy was defined more abstractly by training on a diverse group of nouns differing in animacy (ANIMATE/INANIMATE), even though this training set consisted of data from fewer participants.

Before discussing the two stages and their implications for words in compositional contexts, the single-word training data warrants a brief note. The first stage of phrase-level animacy encoding generalizes from very early on when training on single-word ANIMAL/LOCATION trials, replicating similar timings of other studies decoding word-level animacy (Simanova et al., 2010; Leonardelli et al., 2019; Dirani & Pylkkänen, 2023). Notably, we found a similarly timed cluster when the classifier was trained on single-word ANIMATE/INANIMATE trials. Together, these findings suggest that animacy as an abstract feature is reflected within the first ∼100–150 ms of lexical access and thus available as input for composition shortly after a word is encountered.

### 4.1 Early encoding of composed concepts precedes traditional combinatory operations

Our results reveal a two-stage process for phrase-level animacy encoding, the first of which demonstrates that the product of combinatorial processes—here, phrase-level animacy—is neurally represented after just a few hundred milliseconds. The onset of this effect overlaps in time with several effects associated with combinatorial operations, including conceptual composition in the left ATL (Bemis & Pylkkänen, 2011; Westerlund & Pylkkänen, 2014; Zhang & Pylkkänen, 2015), syntactic processing in the left PTL (Flick et al., 2020; Matchin et al., 2019; Matar et al., 2021), and, most importantly considering our spatial decoding results, the left vmPFC, which has been implicated in prior studies on complement coercion (Pylkkänen & McElree, 2007; Pylkkänen et al., 2009; Brennan & Pylkkänen, 2008, 2010) and hypothesized to represent the output of composition (Pylkkänen, 2019). Crucially, our decoding approach allows us to adjudicate between composition and its representational output in a way that traditional univariate analyses cannot. Because phrase-level animacy in cases like *plastic tiger* is necessarily the result of modifying *tiger* with *plastic,* our ability to decode it provides direct evidence that early neural signals do not merely reflect combinatory operations but also convey the representational content of the composed concept.

How can the output of composition be decoded if combinatory operations are still ongoing? One possibility is that phrase-level animacy is decodable *because* of compositional processes: To integrate words effectively, the brain may maintain relevant semantic features in an active state to constrain the encoding of upcoming words. In the case of *plastic tiger*, early phrase-level decodability could reflect persistent activation of the “inanimate” feature from *plastic* or proactive encoding of the phrase-level concept. Since our stimuli were presented one-by-one and followed a templatic formula, this is an entirely plausible explanation. Future work could thus investigate the exact origins of the inanimate feature by searching for traces of material modifiers within the composed representation (see, e.g., Fyshe et al. (2019) for decoding of adjectives at subsequent nouns). Another possibility, however, is that our results reflect the neural representation of the fully composed concept, rather than the ongoing modulation of discrete semantic features. If such representations are available after a few hundred milliseconds, what has hitherto been construed as indices of combinatory operations in the left ATL, PTL, and vmPFC may instead reflect the cross-cortical journey of a composed concept.

While our decoding approach shows early encoding of phrase-level features, it— like traditional univariate analyses—does not allow us to ascertain whether this reflects compositional processes operating on such features or the final representation of the phrase-level concept. However, an emerging line of research utilizing a stimulus delivery technique flashing short, multiword expressions for a few hundred milliseconds provides independent evidence that early neural signals index the *output* of composition rather than the combinatory operations themselves. Specifically, the brain has been found to exhibit sensitivity to linguistically well-formed expressions as compared to unstructured stimuli beginning at 130 ms post-stimulus onset (Fallon & Pylkkänen, 2024), with responses further being modulated by various linguistic properties (Fallon & Pylkkänen, 2024; Flower & Pylkkänen, 2024, 2026a; Dunagan et al., 2025; Krogh & Pylkkänen, 2025; Li & Pylkkänen, 2026). Crucially, such signals do not scale with the number of operations involved; in fact, the neural signatures remain remarkably stable across one, two, and four words (Flower & Pylkkänen, 2026a), strongly suggesting that they reflect the representation of the entire structure regardless of its size. If the brain can construct and evaluate linguistic structures from simultaneously presented words in just over 100 ms, it seems unlikely that the composition of one word with the preceding one(s), as in our study, should take twice as long.

### 4.2 Sustained representational activity for semantically complex phrases

In the second stage of phrase-level animacy encoding, our temporal decoding analyses revealed a prolonged effect specific to animacy-mismatch phrases. This effect overlapped in time with the late positivity (550–750 ms) for animacy-mismatch phrases relative to controls observed by Schumacher (2013) and further mirrors neural activity deflections reported for other semantically complex expressions like privative adjectives (Schumacher et al., 2018; Fritz & Baggio, 2020), metonymy (Rapp et al., 2011; Schumacher, 2014; Weiland et al., 2014; Weiland-Breckle & Schumacher, 2017; Yurchenko et al., 2020), and complement coercion (Pylkkänen & McElree, 2007; Brennan & Pylkkänen, 2008, 2010; Pylkkänen et al., 2009; Husband et al., 2011; De Almeida et al., 2016; Lai et al., 2017). Moving into source space, our spatial decoding analyses identified the right IFG as one neural generator of the effect. This region has previously been linked to the resolution of metonymic expressions (Rapp et al., 2011) and complement coercion (De Almeida et al., 2016). More broadly, it has been suggested that the right hemisphere as a whole plays a specialized role part in the interpretation of non-literal expressions (e.g., Jung-Beeman, 2005; Johns et al., 2008; Yang, 2014; Diaz & Eppes, 2018; Huang et al., 2023), a category that arguably includes animacy-mismatch phrases. Why might phrase-level animacy remain decodable from the neural signals associated with animacy-mismatch phrases long after it has faded for our controls? Regardless of one’s theoretical stance, computing *plastic tiger* must involve extra “work” relative to *Arabian tiger,* if only because it requires accessing an intuitively less basic—and therefore less frequent—meaning of *tiger*. We observed traces of such processing costs in our behavioral data, with delayed responses for animacy-mismatch phrases in mismatch trials. Based on our temporal decoding results, there are at least two potential drivers for this sustained effect: It may reflect the active maintenance of the “inanimate” feature while combinatory operations act upon constituent words, or it may reflect sustained maintenance of the unified, phrase-level representation. A feature maintenance account is particularly compelling if one assumes a shift from a basic meaning to a more derived one; however, our inability to decode word-level animacy for animacy-mismatch phrases provides little support for such an interpretation. Turning to the alternative explanation, namely sustained animacy maintenance at the level of the phrase, an obvious question is why this would only occur for animacy-mismatch phrases and not our controls. While our data do not shed light on this, one speculative hypothesis is that extra computational efforts result in increased residual activation. This would explain why the inanimacy of animacy-mismatch phrases remains detectable in the neural signals significantly longer than for our controls.

### 4.3 Underspecified lexical representations enriched by context

That phrase-level animacy can override word-level animacy is striking. How can it be that *tiger* in *plastic tiger* is neurally similar to locations (*valley, forest, ocean,* etc.) or inanimates broadly construed (*guitar, blouse, canoe,* etc.) but not to itself and other animals? These results can be accommodated under the underspecification hypothesis, according to which a word’s lexical representation is constrained by the construction in which it appears. In our animacy-mismatch phrases, the notion of *tiger* as a living, breathing animal is barred from participating in the composed concept by the material adjective *plastic*. Conversely, the context of our controls (*Arabian tiger*) imposes no such constraints, allowing the default sentient meaning of *tiger* to emerge.

We found no evidence of word-level animacy encoding for our animacy-mismatch phrases, a result that is irreconcilable with the “first activate, then suppress” trajectory posited by the rich lexical representations hypothesis. Although absence of evidence does not inherently entail evidence of absence, the possibility that word-level effects were simply too weak to detect is unlikely. In fact, word-level effects should be more prominent than phrase-level effects: While the latter relies solely on semantic features arising from composition, the former benefits from identity of both meaning and visual percept across training and test data. That our classifiers successfully decoded the more abstract phrase-level animacy while failing to detect any trace of the perceptually grounded word-level signal strengthens an interpretation of these null results as true evidence of absence. Finally, our results mirror those of a recent decoding study, which also failed to decode word-level properties of nouns embedded in private phrases like *fake salad* while succeeding on subsective phrases like *bad salad* (Law et al., 2026). Altogether, this highlights the impact of context on conceptual representations.

How, then, can the lack of word-level animacy encoding in the current study be reconciled with prior psycholinguistic evidence for the activation of contextually irrelevant features in general (e.g., Swinney, 1979; Glucksberg et al., 2001; Rubio Fernandez, 2007) and, in particular, Hogeweg’s (2019) study using animacy-mismatch phrases? This study employed a Dutch cross-modal priming study with a lexical decision task and found a small but significant priming effect for targets congruent (e.g., *mane*) and incongruent (e.g., *roars*) with prime sentences like *In front of the museum, there is a stone lion.* Crucially, however, this effect may have been driven, or at least impacted, by differences in adjective-noun transition probabilities (e.g., *stone lion* vs. *stone kite*) and the inclusion of representations of artifact-phrases (e.g., *cloth bike* and *felt cigar*). Although we do find examples of artifact representations such as *plastic car* in the real world, the use of material modifiers signals distinct types of semantic shifts for animals versus artifacts:

Whereas *plastic* in *plastic tiger* entails an animate-to-inanimate ontological shift*, plastic* in *plastic car* represents a (marked) departure from a car’s prototypical material, potentially denoting either a toy or a cheaply constructed vehicle. Considering that constitutive materials are routinely used to create inanimate representations of living things—think again of rubber ducks, chocolate bunnies, and the marble lions in front of the New York Public Library—the brain likely leverages these construction-level cues to bypass irrelevant features and directly access the contextually relevant sense of the word in ways not possible for less productive patterns (Klein & Murphy, 2001; Murphy, 2002).

## 5 CONCLUSIONS

Neighboring words may critically shape lexical representations, allowing the most basic word meanings to be divorced from the meaning of a composed concept. Here, we examined one such case, probing the neural encoding of *tiger* in animacy-mismatch phrases like *plastic tiger* to see if the tiger-as-an-animal meaning is retained when it conflicts with the animacy of the overall phrase. While our temporal decoding analyses provided no independent evidence for neural encoding of word-level animacy, we observed a two-stage encoding of phrase-level animacy. Early neural signals recruiting, among other regions, left ventromedial prefrontal cortex, contained reflexes of phrase-level animacy regardless of experimental condition. This was followed by a later, prolonged effect specific to animacy-mismatch phrases that had origins in, among other regions, the right inferior frontal gyrus. The absence of word-level animacy encoding effects for animacy-mismatch phrases counters theories assuming rich lexical representations, which necessitate a shift from one sense of a word to another. Rather, our results suggest that compositional context either enriches underspecified lexical representations or facilitates direct access to contextually relevant senses.

## ACKNOWLEDGEMENTS

This work was supported by the National Science Foundation award #2335767 (LP). We thank Bernarda Basualdo for assistance with data collection as well as Alec Marantz, Ailís Cournane, Sebastian Michelmann, Alona Fyshe, and the members of NeLLab for their suggestions and feedback.

## COMPETING INTERESTS

The authors declare no competing interests.

## AUTHOR CONTRIBUTIONS

Conceptualization: SK, LP

Methodology: SK, LP

Investigation: SK

Data curation: SK

Formal analysis: SK

Software: SK

Resources: SK

Validation: SK, LP

Visualization: SK

Funding acquisition: LP

Project administration: SK, LP

Supervision: LP

Writing – original draft: SK

Writing – review & editing: SK, LP

## DECLARATION OF GENERATIVE AI AND AI-ASSISTED TECHNOLOGIES IN THE MANUSCRIPT PREPARATION PROCESS

During the preparation of this work the authors used Cursor, an AI-assisted code editor, for limited code writing, editing, and debugging of existing analysis scripts, using both automatic model routing (Auto) and manually selected Claude Sonnet and Opus models (4.5 and 4.6). Google Gemini was used for minor editing in the final writing stages. After using these tools, the authors reviewed and edited the content as needed and take full responsibility for the content of the published article.

## Notes

https://osf.io/bgkfm/overview

### Competing Interest Statement

The authors have declared no competing interest.

## REFERENCES

Adachi, Y., Shimogawara, M., Higuchi, M., Haruta, Y., & Ochiai, M. (2001). Reduction of non-periodic environmental magnetic noise in MEG measurement by Continuously Adjusted Least Squares Method. IEEE Transactions on Applied Superconductivity, 11, 669–672.

Asher, N. (2011). Lexical meaning in context: A web of words (1st ed.). Cambridge University Press. 10.1017/CBO9780511793936

Asher, N., Abrusan, M., & Van De Cruys, T. (2017). Types, meanings and co-composition in lexical semantics. In S. Chatzikyriakidis & Z. Luo (Eds.), Modern Perspectives in Type-Theoretical Semantics (Vol. 98, pp. 135–161). Springer International Publishing. 10.1007/978-3-319-50422-3_6

Balota, D. A., Yap, M. J., Cortese, M. J., Hutchison, K. A., Kessler, B., Loftis, B., Neely, J. H., Nelson, D. L., Simpson, G. B., & Treiman, R. (2007). The English Lexicon Project. Behavior Research Methods, 39(3), 445–459.

Bates, D., Mächler, M., Bolker, B., & Walker, S. (2015). Fitting linear mixed-effects models using lme4. Journal of Statistical Software, 67(1). 10.18637/jss.v067.i01

Bemis, D. K., & Pylkkänen, L. (2011). Simple composition: A magnetoencephalography investigation into the comprehension of minimal linguistic phrases. The Journal of Neuroscience, 31(8), 2801–2814. 10.1523/JNEUROSCI.5003-10.2011

Benjamini, Y., & Hochberg, Y. (1995). Controlling the false discovery rate: A practical and powerful approach to multiple testing. Journal of the Royal Statistical Society: Series B (Methodological), 57(1), 289–300. 10.1111/j.2517-6161.1995.tb02031.x

Bierwisch, M., & Schreuder, R. (1992). From concepts to lexical items. Cognition, 42(1– 3), 23–60. 10.1016/0010-0277(92)90039-K

Blutner, R. (1998). Lexical pragmatics. Journal of Semantics, 15(2), 115–162. 10.1093/jos/15.2.115

Blutner, R. (2004). Pragmatics and the lexicon. In L. R. Horn & G. Ward (Eds.), The handbook of pragmatics (pp. 488–514). Wiley Online Library.

Brennan, J., & Pylkkänen, L. (2008). Processing events: Behavioral and neuromagnetic correlates of aspectual coercion. Brain and Language, 106(2), 132–143. 10.1016/j.bandl.2008.04.003

Brennan, J., & Pylkkänen, L. (2010). Processing psych verbs: Behavioural and MEG measures of two different types of semantic complexity. Language and Cognitive Processes, 25(6), 777–807. 10.1080/01690961003616840

Brysbaert, M., & New, B. (2009). Moving beyond Kučera and Francis: A critical evaluation of current word frequency norms and the introduction of a new and improved word frequency measure for American English. Behavior Research Methods, 41(4), 977–990. 10.3758/BRM.41.4.977

Carston, R. (2012). Word meaning and concept expressed. The Linguistic Review, 29(4), 607–623.

Dale, A. M., Liu, A. K., Fischl, B. R., Buckner, R. L., Belliveau, J. W., Lewine, J. D., & Halgren, E. (2000). Dynamic Statistical Parametric Mapping. Neuron, 26(1), 55– 67. 10.1016/S0896-6273(00)81138-1

Davies, M. (2008). The Corpus of Contemporary American English (COCA): One billion words, 1990-2019 [Computer software]. https://www.english-corpora.org/coca/

Davies, M. (2018). The iWeb Corpus [Computer software]. https://www.english-corpora.org/iWeb/

De Almeida, R. G., Riven, L., Manouilidou, C., Lungu, O., Dwivedi, V. D., Jarema, G., & Gillon, B. (2016). The neuronal correlates of indeterminate sentence comprehension: An fMRI study. Frontiers in Human Neuroscience, 10. 10.3389/fnhum.2016.00614

Del Pinal, G. (2015). Dual Content Semantics, privative adjectives, and dynamic compositionality. Semantics and Pragmatics, 8. 10.3765/sp.8.7

Desikan, R. S., Ségonne, F., Fischl, B., Quinn, B. T., Dickerson, B. C., Blacker, D., Buckner, R. L., Dale, A. M., Maguire, R. P., Hyman, B. T., Albert, M. S., & Killiany, R. J. (2006). An automated labeling system for subdividing the human cerebral cortex on MRI scans into gyral based regions of interest. NeuroImage, 31(3), 968–980. 10.1016/j.neuroimage.2006.01.021

Diaz, M. T., & Eppes, A. (2018). Factors influencing right hemisphere engagement during metaphor comprehension. Frontiers in Psychology, 9, 414. 10.3389/fpsyg.2018.00414

Dirani, J., & Pylkkänen, L. (2023). The time course of cross-modal representations of conceptual categories. NeuroImage, 277, 120254. 10.1016/j.neuroimage.2023.120254

Dunagan, D., Jordan, T., Hale, J. T., Pylkkänen, L., & Chacón, D. A. (2025). Evaluating the timecourses of morpho-orthographic, lexical, and grammatical processing following rapid parallel visual presentation: An EEG investigation in English. Cognition, 257, 106080. 10.1016/j.cognition.2025.106080

Fallon, J., & Pylkkänen, L. (2024). Language at a glance: How our brains grasp linguistic structure from parallel visual input. Science Advances, 10(43), eadr9951. 10.1126/sciadv.adr9951

Fischl, B. (2012). FreeSurfer. NeuroImage, 62(2), 774–781. 10.1016/j.neuroimage.2012.01.021

Flick, G., & Pylkkänen, L. (2020). Isolating syntax in natural language: MEG evidence for an early contribution of left posterior temporal cortex. Cortex, 127, 42–57. 10.1016/j.cortex.2020.01.025

Flower, N., & Pylkkänen, L. (2024). The spatiotemporal dynamics of bottom-up and top-down processing during at-a-glance reading. The Journal of Neuroscience, 44(48), e0374242024. 10.1523/JNEUROSCI.0374-24.2024

Flower, N., & Pylkkänen, L. (2026a). A unified neural time course for words, phrases, and sentences: MEG evidence from parallel presentation. Neurobiology of Language, 1–26. 10.1162/NOL.a.254

Flower, N., & Pylkkänen, L. (2026b). Cortical representation of quantification: The role of the left anterior temporal lobe. Cognition, 270, 106403. 10.1016/j.cognition.2025.106403

Frisson, S. (2009). Semantic underspecification in language processing. Language and Linguistics Compass, 3(1), 111–127. 10.1111/j.1749-818X.2008.00104.x

Fritz, I., & Baggio, G. (2020). Meaning composition in minimal phrasal contexts: Distinct ERP effects of intensionality and denotation. Language, Cognition and Neuroscience, 35(10), 1295–1313. 10.1080/23273798.2020.1749678

Fyshe, A., Sudre, G., Wehbe, L., Rafidi, N., & Mitchell, T. M. (2019). The lexical semantics of adjective–noun phrases in the human brain. Human Brain Mapping, 40(15), 4457–4469. 10.1002/hbm.24714

Glucksberg, S., Newsome, M. R., & Goldvarg, Y. (2001). Inhibition of the literal: Filtering metaphor-irrelevant information during metaphor comprehension. In Models of figurative language (pp. 277–293). Psychology Press.

Gramfort, A., Luessi, M., Larson, E., Engemann, D. A., Strohmeier, D., Brodbeck, C., Parkkonen, L., & Hämäläinen, M. S. (2014). MNE software for processing MEG and EEG data. NeuroImage, 86, 446–460. 10.1016/j.neuroimage.2013.10.027

Hogeweg, L. (2019). Suppression in interpreting adjective noun combinations and the nature of the lexicon. Journal of Semantics, 36(4), 721–751. 10.1093/jos/ffz012

Hogeweg, L., & Vicente, A. (2020). On the nature of the lexicon: The status of rich lexical meanings. Journal of Linguistics, 56(4), 865–891. 10.1017/S0022226720000316

Huang, Y., Huang, J., Li, L., Lin, T., & Zou, L. (2023). Neural network of metaphor comprehension: An ALE meta-analysis and MACM analysis. Cerebral Cortex, bhad337. 10.1093/cercor/bhad337

Husband, E. M., Kelly, L. A., & Zhu, D. C. (2011). Using complement coercion to understand the neural basis of semantic composition: Evidence from an fMRI study. Journal of Cognitive Neuroscience, 23(11), 3254–3266. 10.1162/jocn_a_00040

Jackendoff, R. (1997). The architecture of language faculty. the MIT press.

Johns, C. L., Tooley, K. M., & Traxler, M. J. (2008). Discourse impairments following right hemisphere brain damage: A critical review. Language and Linguistics Compass, 2(6), 1038–1062. 10.1111/j.1749-818X.2008.00094.x

Jung-Beeman, M. (2005). Bilateral brain processes for comprehending natural language. Trends in Cognitive Sciences, 9(11), 512–518. 10.1016/j.tics.2005.09.009

King, J.-R., & Dehaene, S. (2014). Characterizing the dynamics of mental representations: The temporal generalization method. Trends in Cognitive Sciences, 18(4), 203–210. 10.1016/j.tics.2014.01.002

Klein, D. E., & Murphy, G. L. (2001). The representation of polysemous words. Journal of Memory and Language, 45(2), 259–282. 10.1006/jmla.2001.2779

Krogh, S., & Pylkkänen, L. (2025). Manipulating syntax without taxing working memory: MEG correlates of syntactic dependencies in a verb-second language. *Language*, Cognition and Neuroscience, 1–24. 10.1080/23273798.2025.2549342

Lai, Y.-Y., Lacadie, C., Constable, T., Deo, A., & Piñango, M. M. (2017). Complement coercion as the processing of aspectual verbs: Evidence from self-paced reading and fMRI. In J. A. Hampton & Y. Winter (Eds.), Compositionality and Concepts in Linguistics and Psychology (Vol. 3, pp. 191–222). Springer International Publishing. 10.1007/978-3-319-45977-6_8

Lambon Ralph, M. A., Lowe, C., & Rogers, T. T. (2006). Neural basis of category-specific semantic deficits for living things: Evidence from semantic dementia, HSVE and a neural network model. Brain, 130(4), 1127–1137. 10.1093/brain/awm025

Law, R. M. C., Lambon Ralph, M. A., & Hauk, O. (2026). A common framework for semantic memory and semantic composition. Imaging Neuroscience, 4, IMAG.a.1131. 10.1162/IMAG.a.1131

Leonardelli, E., Fait, E., & Fairhall, S. L. (2019). Temporal dynamics of access to amodal representations of category-level conceptual information. Scientific Reports, 9(1), 239. 10.1038/s41598-018-37429-2

Li, B., & Pylkkänen, L. (2026). MEG investigation of adjective order preferences as a syntactic constraint. Cognition, 273, 106536. 10.1016/j.cognition.2026.106536

Maris, E., & Oostenveld, R. (2007). Nonparametric statistical testing of EEG- and MEG-data. Journal of Neuroscience Methods, 164(1), 177–190. 10.1016/j.jneumeth.2007.03.024

Matar, S., Dirani, J., Marantz, A., & Pylkkänen, L. (2021). Left posterior temporal cortex is sensitive to syntax within conceptually matched Arabic expressions. Scientific Reports, 11(1), 7181. 10.1038/s41598-021-86474-x

Matchin, W., Brodbeck, C., Hammerly, C., & Lau, E. (2019). The temporal dynamics of structure and content in sentence comprehension: Evidence from fMRI-constrained MEG. Human Brain Mapping, 40(2), 663–678. 10.1002/hbm.24403

Murphy, G. L. (2002). Word meaning. In The Big Book of Concepts (pp. 385–441). The MIT Press. 10.7551/mitpress/1602.001.0001

Partee, B. H. (2010). Privative adjectives: Subsective plus coercion. In R. Bäuele, U. Reyle, & T. E. Zimmermann (Eds.), Presuppositions and discourse: Essays offered to Hans Kamp (pp. 273–285). Emerald Group Publishing.

Pedregosa, F., Varoquaux, G., Gramfort, A., Michel, V., Thirion, B., Grisel, O., Blondel, M., Prettenhofer, P., Weiss, R., Dubourg, V., Vanderplas, J., Passos, A., Cournapeau, D., Brucher, M., Perrot, M., & Duchesnay, E. (2011). Scikit-learn: Machine learning in Python. Journal of Machine Learning Research, 12, 2825–2830.

Pegado, F., Wen, Y., Mirault, J., Dufau, S., & Grainger, J. (2021). An ERP investigation of transposed-word effects in same-different matching. Neuropsychologia, 153, 107753. 10.1016/j.neuropsychologia.2021.107753

Piñango, M. M., Zhang, M., Foster-Hanson, E., Negishi, M., Lacadie, C., & Constable, R. T. (2017). Metonymy as referential dependency: Psycholinguistic and neurolinguistic arguments for a unified linguistic treatment. Cognitive Science, 41(S2), 351–378. 10.1111/cogs.12341

Pustejovsky, J. (1995). The Generative Lexicon. The MIT Press. 10.7551/mitpress/3225.001.0001

Pylkkänen, L. (2019). The neural basis of combinatory syntax and semantics. Science, 366(6461), 62–66. 10.1126/science.aax0050

Pylkkänen, L., Martin, A. E., McElree, B., & Smart, A. (2009). The Anterior Midline Field: Coercion or decision making? Brain and Language, 108(3), 184–190. 10.1016/j.bandl.2008.06.006

Pylkkänen, L., & McElree, B. (2007). An MEG study of silent meaning. Journal of Cognitive Neuroscience, 19(11), 1905–1921. 10.1162/jocn.2007.19.11.1905

Ralph, M. A. L., Jefferies, E., Patterson, K., & Rogers, T. T. (2017). The neural and computational bases of semantic cognition. Nature Reviews Neuroscience, 18(1), 42–55. 10.1038/nrn.2016.150

Rapp, A. M., Erb, M., Grodd, W., Bartels, M., & Markert, K. (2011). Neural correlates of metonymy resolution. Brain and Language, 119(3), 196–205. 10.1016/j.bandl.2011.07.004

Reyle, U. (1993). Dealing with ambiguities by underspecification: Construction, representation and deduction. Journal of Semantics, 10(2), 123–179. 10.1093/jos/10.2.123

Rubio Fernandez, P. (2007). Suppression in metaphor interpretation: Differences between meaning selection and meaning construction. Journal of Semantics, 24(4), 345–371. 10.1093/jos/ffm006

Sassenhagen, J., & Draschkow, D. (2019). Cluster-based permutation tests of MEG/EEG data do not establish significance of effect latency or location. Psychophysiology, 56(6), e13335. 10.1111/psyp.13335

Schumacher, P. B. (2013). When combinatorial processing results in reconceptualization: Toward a new approach of compositionality. Frontiers in Psychology, 4. 10.3389/fpsyg.2013.00677

Schumacher, P. B. (2014). Content and context in incremental processing: “The ham sandwich” revisited. Philosophical Studies, 168(1), 151–165. 10.1007/s11098-013-0179-6

Schumacher, P. B., Brandt, P., & Weiland-Breckle, H. (2018). Online processing of “real” and “fake”: The cost of being too strong. In E. Castroviejo, L. McNally, & G. Weidman Sassoon (Eds.), The Semantics of Gradability, Vagueness, and Scale Structure (Vol. 4, pp. 93–111). Springer International Publishing. 10.1007/978-3-319-77791-7_4

Simanova, I., Van Gerven, M., Oostenveld, R., & Hagoort, P. (2010). Identifying object categories from event-related EEG: Toward decoding of conceptual representations. PLoS ONE, 5(12), e14465. 10.1371/journal.pone.0014465

Singmann, H., Bolker, B., Westfall, J., Aust, F., & Ben-Schachar, M. (2023). afex: Analysis of Factorial Experiments (Version R package version 1.3-0) [Computer software]. https://CRAN.R-project.org/package=afex

Swinney, D. A. (1979). Lexical access during sentence comprehension: (Re)consideration of context effects. Journal of Verbal Learning and Verbal Behavior, 18(6), 645–659. 10.1016/S0022-5371(79)90355-4

Vicente, A. (2015). The green leaves and the expert: Polysemy and truth-conditional variability. Lingua, 157, 54–65. 10.1016/j.lingua.2014.04.013

Weiland, H., Bambini, V., & Schumacher, P. B. (2014). The role of literal meaning in figurative language comprehension: Evidence from masked priming ERP. Frontiers in Human Neuroscience, 8. 10.3389/fnhum.2014.00583

Weiland-Breckle, H., & Schumacher, P. B. (2017). Artist-for-work metonymy: Type clash or underspecification? The Mental Lexicon, 12(2), 219–233. 10.1075/ml.16014.wei

Westerlund, M., & Pylkkänen, L. (2014). The role of the left anterior temporal lobe in semantic composition vs. Semantic memory. Neuropsychologia, 57, 59–70. 10.1016/j.neuropsychologia.2014.03.001

Wisniewski, E. J. (1996). Construal and similarity in conceptual combination. Journal of Memory and Language, 35(3), 434–453. 10.1006/jmla.1996.0024

Yang, J. (2014). The role of the right hemisphere in metaphor comprehension: A metaanalysis of functional magnetic resonance imaging studies: Right hemisphere in metaphor processing. Human Brain Mapping, 35(1), 107–122. 10.1002/hbm.22160

Yurchenko, A., Lopukhina, A., & Dragoy, O. (2020). Metaphor is between metonymy and homonymy: Evidence from event-related potentials. Frontiers in Psychology, 11, 2113. 10.3389/fpsyg.2020.02113

Zhang, L., & Pylkkänen, L. (2015). The interplay of composition and concept specificity in the left anterior temporal lobe: An MEG study. NeuroImage, 111, 228–240. 10.1016/j.neuroimage.2015.02.028

